# Genomic epidemiology of ESBL-producing *Escherichia coli* from humans and an Aotearoa New Zealand river

**DOI:** 10.1101/2024.05.05.592593

**Authors:** Holly A Gray, Patrick J Biggs, Anne C Midwinter, Lynn E Rogers, Ahmed Fayaz, Rukhshana N Akhter, Sara A Burgess

## Abstract

In Aotearoa New Zealand, urinary tract infections in humans are commonly caused by extended-spectrum beta-lactamase (ESBL)-producing *Escherichia coli*. This group of antimicrobial resistant bacteria are often multidrug resistant. However, there is limited information on ESBL-producing *E. coli* found in the environment and their link with human clinical isolates. In this study, we examined the genetic relationship of environmental and human clinical ESBL-producing *E. coli* and isolates collected in parallel within the same area over 14 months. Environmental samples were collected from treated effluent, stormwater and multiple locations along an Aotearoa New Zealand river. Treated effluent, stormwater and river water sourced downstream of the treated outflow point were the main sources of ESBL-producing *E. coli* (7/14 samples, 50.0%; 3/6 samples, 50%; and 15/28 samples, 54% respectively). Whole genome sequence comparison was carried out on 307 human clinical and 45 environmental ESBL-producing *E. coli* isolates. Sequence type 131 was dominant for both clinical (147/307, 47.9%) and environmental isolates (11/45, 24.4%). The most prevalent ESBL genes were both *bla*_CTX-M-27_ and *bla*_CTX-M-15_ for the clinical isolates (134/307, 43.6%) and *bla*_CTX-M-15_ for the environmental isolates (28/45, 62.2%). A core single nucleotide polymorphism analysis of these isolates suggested that some strains were shared between humans and the local river. These results highlight the importance of understanding different transmission pathways for the spread of ESBL-producing *E. coli*.

**2. Impact statement:** Extended spectrum beta lactamase (ESBL)-producing *E. coli* frequently cause urinary tract infections that exhibit multidrug resistance. Surveillance studies have identified the predominant strains and resistance genes associated with urinary tract infections. However, there is limited information on the extent of spread beyond the patient. We describe the genetic relatedness of ESBL-producing environmental and clinical *E. coli* isolated during the same temporal-spatial period in Aotearoa New Zealand. Comparative genomic analyses of these bacteria provide evidence of clonal spread between humans and the environment, highlighting the need to integrate environmental surveillance into antimicrobial resistance monitoring.

**3. Data summary:** All Illumina sequence reads for this study have been deposited in GenBank under BioProject PRJNA1032159, except for strain SB0283h1, whose data can be found under BioProject PRJNA715472. The sequence read accessions for each genome are provided in the supplementary material.

The code used for the genomic and statistical analyses is available from the GitHub repository https://github.com/sburgess1/Manawat-_ESBL.

**The authors confirm all supporting data and protocols have been provided within the article or through supplementary data files.**

## 4 Introduction

*Escherichia coli* are gram-negative bacteria that form a natural part of the mammalian intestinal tract microbiota. They can be opportunistic pathogens and can cause a range of infections in humans such as pneumonia, bacteremia, meningitis, and urinary tract infections [1–3]. The spread of antimicrobial resistance in pathogenic strains of *E. coli* is of importance because infections caused by these strains are often harder to treat, resulting in increased severity and duration of infection [4, 5]. One important group of antimicrobial resistant *E. coli* is the extended-spectrum beta-lactamase (ESBL)-producing *E. coli*. In Aotearoa New Zealand (NZ), *E. coli* was associated with 84.6 % of ESBL-producing Enterobacterale*s* related infections in 2019 [6].

The ESBL enzymes confer resistance to an extended range of beta-lactam antibiotics, including the first, third and some fourth generation cephalosporins but not the cephamycins or carbapenems [7, 8]. CTX-M is the predominant enzyme type associated with human clinical ESBL-producing *E. coli* [9–11]. There are more than 250 *bla*_CTX-M_ gene variants, with *bla*_CTX-M-15_ being the most common amongst human clinical *E. coli* [12–15]. The genes encoding ESBLs are often associated with other antimicrobial resistance genes (ARGs), resulting in a multidrug resistant (MDR) strain [10, 11, 16]. MDR pathogenic strains of *Enterobacteriaceae* are of particular concern to the health sector resulting in ESBL-producing *Enterobacteriaceae* being identified as “critical” on the World Health Organisation’s “Priority Pathogens List” [17].

The main route for community dissemination of antimicrobial-resistant bacteria involves person-to-person transmission [18, 19]. However, there are alternative transmission pathways, such as contact with companion and livestock animals, consumption of food products, or indirect exposure through contaminated recreational water [20–23]. The natural environment and particularly recreational water have been identified as important vectors in the transmission of antimicrobial resistant bacteria [24, 25]. Recreational water use has rarely been found to be a risk factor for ESBL infection and the carriage of ESBL-producing *E. coli* in recreational water users has not been found to be higher compared to non-recreational water users [26–29]. However, freshwater has been identified as a source of ESBL-producing *E. coli* [30, 31]. There are few studies that have compared ESBL-producing *E. coli* from human clinical samples and freshwater samples [32–34]. The aim of this study was to compare the genetic relatedness of clinical and environmental ESBL-producing *E. coli*. Whole genome sequence analysis was carried out for isolates collected over the same time period and geographical region.

## 5 Methods

### 5.1 Sample collection and processing

Environmental samples were collected from the Manawatū River (sites A, B, D and F), stormwater (site C), and treated effluent (site E) over 14 months: August 2019 to March 2020 and July 2020 to January 2021 (excluding October 2019). Two water samples and one sediment sample were collected from locations A (upstream of the city of Palmerston North), B, D, and F (downstream) along the Manawatū River (Figure 1). A stormwater sample was collected from the Centennial Drive site (site C) when the stormwater flowed from the drain.

**Figure 1:**
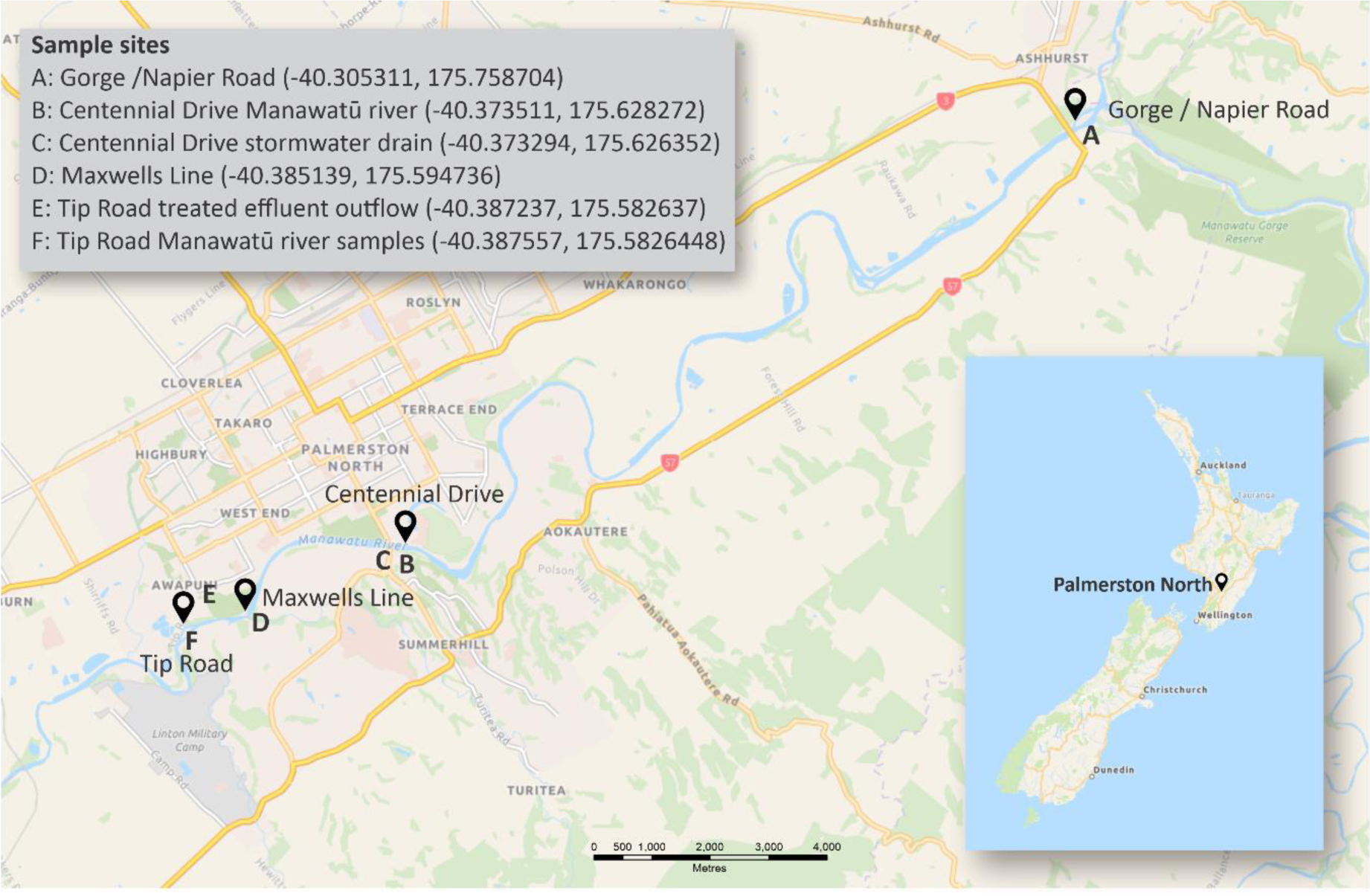
Sampling sites A to F along the Manawatū river, located in Palmerston North Aotearoa New Zealand. River flow is the direction of site A to F. Image produced using CorelDRAW® 2019 (Corel Corporation). Base map obtained from Google Maps (2023), available at: https://www.google.com/maps/@-40.3521071,175.6424104,12.87z?entry=ttu (Accessed: 16 August 2023).

Water samples prepared as previously described [35] and sediment (1 g) were enriched in 10 ml of buffered peptone water (BPW; BD Difco^TM^, Becton Dickinson) or *E. coli* broth (ECB; Oxoid Ltd) and incubated overnight at 35°C. Additionally, 1 ml of treated effluent was diluted in 9 ml of BPW and/or ECB and incubated overnight at 35°C. Two millilitres of each enrichment was centrifuged for 5 min at 6000 g and the pellet was resuspended in 1ml of in-house prepared nutrient broth no. 2 (Oxoid Ltd) containing 15% (v/v) glycerol then stored at -80°C. Each sample was allocated a unique identifying number with the prefix “SB” followed by the allocated sample number. The enrichments were plated onto two selective MacConkey agar plates (containing 1 µg/ml of cefotaxime sodium or ceftazidime) and CHROMagar™ ESBL (Fort Richard Laboratories) as previously described [36]. Additionally, 1 ml of treated effluent was plated onto CHROMagar™ ESBL. After incubation at 35°C for 16 to 24 hours, presumptive *E. coli* colonies (up to two colonies from each plate) were purified on Columbia horse blood agar (Fort Richard Laboratories) and identified using matrix-assisted laser desorption ionization-time of flight (MALDI-TOF Biotyper®, Bruker Daltonics) [36, 37]. Presumptive ESBL-producing *Enterobacteriaceae* clinical strains, which originated from urine, blood, or wound swab cultures were isolated by MedLab Central Laboratory (Palmerston North, Aotearoa New Zealand), from August 2019 to March 2020 and June 2020 to January 2021. These isolates were also purified on Columbia horse blood agar, identified using MALDI-TOF mass spectrometry, and allocated a unique identifying number with the prefix “EH”.

### 5.2 Antimicrobial susceptibility testing

Antimicrobial susceptibility tests and confirmation of an ESBL-producing phenotype were carried out using Kirby-Bauer disc diffusion assays following Clinical and Laboratory Standards Institute guidelines [38]. Antimicrobial susceptibility tests of the environmental *E. coli* isolates were carried out using a panel of ten antibiotics (MAST Group, Table S1). ESBL confirmation was carried out on the clinical *E. coli* as well as those environmental *E. coli* that were non-susceptible to cefotaxime or ceftazidime using the double-disc comparison assay (D62C cefotaxime and D64C ceftazidime ESBL disc tests; Mast Group).

### 5.3 DNA Extractions and whole genome sequencing

ESBL-producing *E. coli* isolates were grown in Luria-Bertani broth (made with deionised water, NaCl (Scharlau), yeast extract (Becton Dickinson Difco^TM^) and tryptone (Becton Dickinson Difco^TM,^)) overnight at 35°C and genomic DNA was extracted using the Wizard® Genomic DNA purification kit (Promega) as previously described [35]. Libraries were generated using the Nextera XT DNA library preparation kit (Illumina Inc) and submitted to the Massey Genome Service (Massey University) for quality checks and sequencing. Whole genome sequencing was performed by Novogene ( Singapore) using Illumina HiSeq™ X sequencing (2 x 150 bp paired-end reads). FastQC (v.0.11.9) was used for the quality assessment of the raw reads [39].

### 5.4 Genome assemblies and analyses

The raw reads were processed using Nullarbor 2 (v.2.0.20191013), using the default parameters and strain *E. coli* EC968 was used as the reference (GenBank accession HG941718.1) [40, 41]. Using this pipeline, the adapters were removed using Trimmomatic (v.0.39), the reads were assembled using SKESA (v.2.4.0) and the assemblies were annotated using Prokka (v. 1.14.6) [42–44]. Quast (v.5.0.2) was used for genome evaluations [45]. Additionally, multi-locus sequence-typing (MLST) was carried out using mlst (v.2.19.0), and resistance and virulence genes were identified using ABRicate (v.1.0.1) with the NCBI database (v.2021-03-27) and VirulenceFinder (v.2021-03-27) databases respectively. The presence of two or more genes as described by [46] (listed in Table S2) was used to determine whether isolates could be classified as extra-intestinal pathogenic *E. coli* (ExPEC) [46]. PointFinder (v.3.0) was used to detect point mutations associated with antimicrobial resistance in the sequences of the ST131 isolates [47]. The *fimH* type was determined using FimTyper (v.1.0) with default parameters for the isolates EH0318a, EH0378a, EH0389a, and EH0391a [48].

ChewBBACA (v.3.1.2) was used to generate a core-genome MLST (cgMLST) allele profile using a 95% threshold [49], and a distance matrix was generated using cgMLST-dists (v.0.4.0, https://github.com/tseemann/cgmlst-dists) [49]. A neighbour-joining tree was generated using the R package ape (v.5.7-1) [50]. A core SNP analysis was carried out using Snippy-multi (v.4.6.0), employing the internal references EH0395a (ST131), EH0143a (ST69), and EH0294a (ST1193) [51]. Gubbins (v.2.3.1) was applied to all Snippy analyses, with the FastTree (v2.1.11) general time-reversible (GTR) substitution model being used for phylogenetic tree construction [52]. Default parameters for both Gubbins and FastTree were used. Phylogenetic trees were uploaded to the Interactive Tree Of Life (iTOL, v.6.5.7) for annotation and visualisation [50]. To classify the ST131 isolates into clades, sequence reads of known ST131 clades A, B and C were downloaded from the European Nucleotide Archive: strains MER-56 (SRR5936479), MER-53 (SRR5936492) and MER-25 (SRR5936501) respectively [14].

### 5.5 Plasmidome analysis

Presumptive plasmid contigs were identified using RFPlasmid (v. 0.0.18) [53] and extracted using a custom Python script (v.3.8.1). The plasmid contigs were annotated using Prokka (v. 1.14.6) and the GFF Prokka-generated files were used as the input for a pangenome analysis using Panaroo (v. 1.3.0), which was run in sensitive mode, using the aligner MAFFT, and a core threshold of 0.95 [54, 55]. The gene presence-absence output from Panaroo was uploaded into Panini (https://panini.cgps.group/ Accessed 27 October 2023) and the resulting csv and dot files were uploaded to Microreact (https://microreact.org/ Accessed 27 October 2023) [56, 57].

### 5.6 Graphical data display

Phenotypic results were stored in a Microsoft Access database (Microsoft 365, v16.0.1) and analysed using the software R (v.4.1.2, R computing group). The code used is available from the GitHub repository: https://github.com/sburgess1/Manawat-_ESBL. R packages used included ggplot2 (v.3.3.5), lubridate (v.1.8.0), ComplexUpset (v.1.3.1), ggalluvial (v.0.12.3), and easyalluvial (v.0.3.0).

## 6 Results

### 6.1 The detection of ESBL-producing and multidrug resistant *E. coli* in freshwater, stormwater, and treated sewage

To evaluate the prevalence and persistence of antimicrobial resistant *E. coli* in a waterway, both water and sediment samples were collected from four locations along the Manawatū River over a 14-month period. ESBL-producing *E. coli* were isolated throughout every season, during periods of both low and high rainfall (Figure S1, Table S3). MDR resistance (resistance to three or more classes ) and ESBL-producing *E. coli* were detected from three of the four river sample sites (Table 1), with sample site F (downstream of the treated sewage outlet) having a significantly higher proportion of both ESBL-producing (15/28, 53.6%, 95% CI: 29.8 – 72.2%, p=1.863e-09) and multidrug resistant *E. coli* (11/28, 39.3%, 95% CI: 15.9 - 57.6%, p = 2.078e-06) than the other three sites. To evaluate incoming sources of antimicrobial resistant *E. coli*, samples were also collected from a stormwater drain (site C) and the treated sewage outlet (site E). A similar proportion (p > 0.05) of ESBL-producing *E. coli* and MDR *E. coli* was observed for both sites when compared to sample site F.

**Table 1:** Sample level prevalence of ESBL-producing, antimicrobial and multidrug resistant E. coli.

|  | A |  |  | B |  |  | C | D |  |  | E | F |  |  |
| --- | --- | --- | --- | --- | --- | --- | --- | --- | --- | --- | --- | --- | --- | --- |
| Sample type | Sediment | Water | Total | Sediment | Water | Total | Storm water | Sediment | Water | Total | Treated effluent | Sediment | Water | Total |
| ESBL | 0/12<br>(0.0%) | 1/28<br>(3.6%) | 1/40<br>(2.5%) | 0/13<br>(0.0%) | 1/28<br>(3.6%) | 1/41<br>(2.4%) | **3/6**<br>**(50.0%)** | 0/11<br>(0.0%) | 0/22<br>(0.0%) | 0/33<br>(0.0%) | **7/14**<br>**(50.0%)** | 1/12<br>(8.3%) | **15/28**<br>**(53.6%)** | 16/40<br>(40.0%) |
| Antimicrobial resistant <sup>a</sup> | 2/12<br>(16.7%) | 2/28<br>(7.1%) | 2/40<br>(5.0%) | 2/13<br>(15.4%) | 2/28<br>(7.1%) | 4/41<br>(9.8%) | **4/6**<br>**(66.7%)** | 0/11<br>(0.0%) | 0/22<br>(0.0%) | 0/33<br>(0.0%) | **7/14**<br>**(50.0%)** | 2/12<br>(16.7%) | **17/28**<br>**(60.7%)** | 19/40<br>(47.5%) |
| Multidrug resistant | 0/12<br>(0.0%) | 1/28<br>(3.6%) | 1/40<br>(2.5%) | 0/13<br>(0.0%) | 1/28<br>(3.6%) | 1/41<br>(2.4%) | 2/6<br>(33.3%) | 0/11<br>(0.0%) | 0/22<br>(0.0%) | 0/33<br>(0.0%) | 6/14<br>(42.9%) | 1/12<br>(8.3%) | 11/28<br>(39.3%) | 12/40<br>(30.0%) |

<sup>a</sup> Resistant to at least one of the ten antibiotics used to screen the samples.

The environmental *E. coli* isolated from the six sample sites were screened for resistance to ten antibiotics (Figure 2). The most observed resistance phenotypes were to cefotaxime, ceftazidime and cefoxitin as well as cefotaxime and ceftazidime. Resistance to trimethoprim-sulfamethoxazole, which is commonly used for the treatment of UTIs [58], was observed in combination with resistance to one or more other antimicrobials. A multidrug resistance phenotype was observed in 35/82 (42.7%) isolates.

**Figure 2.**
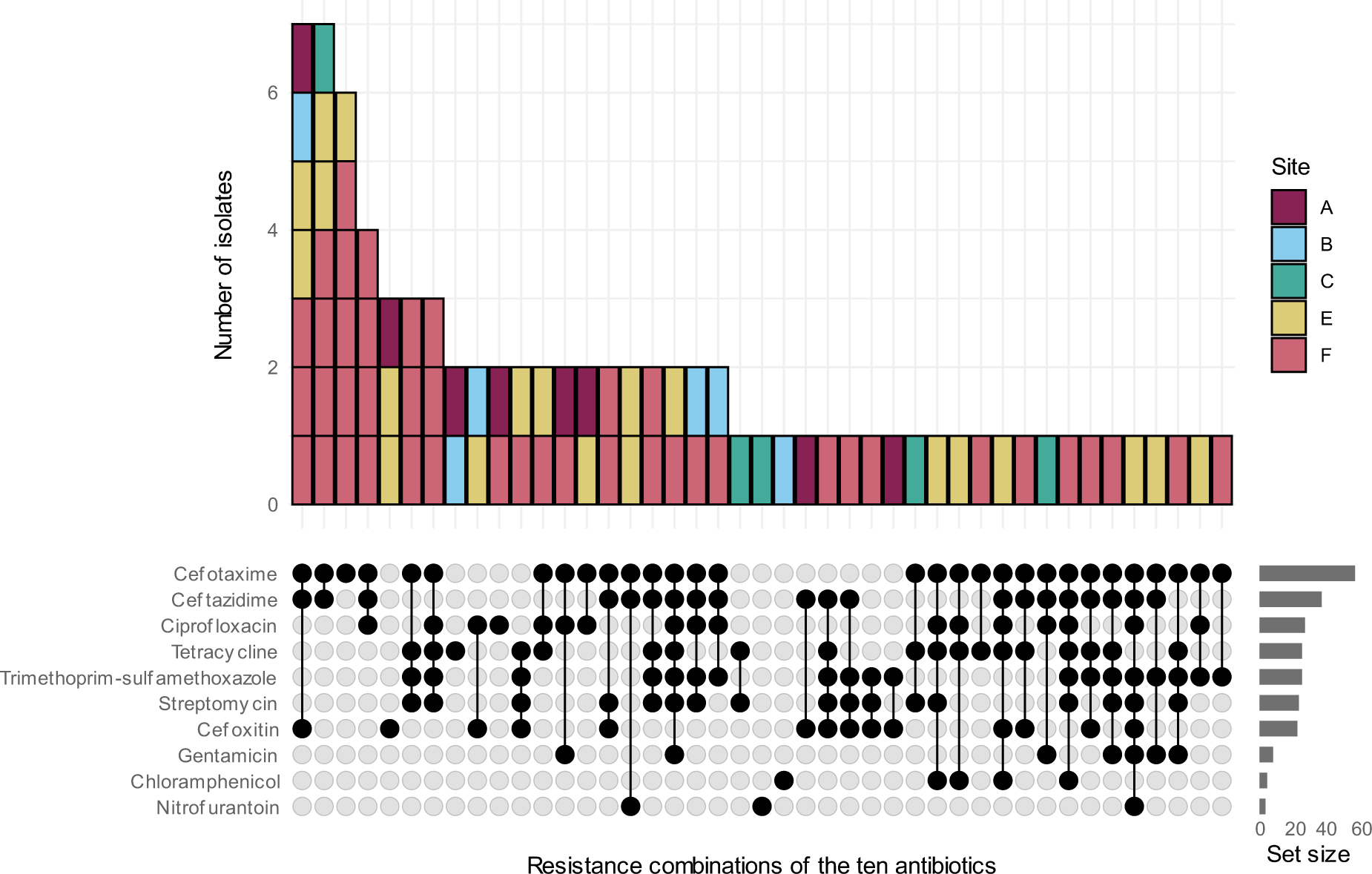
Antimicrobial resistance profiles of ESBL-producing *E. coli* against ten antibiotics from sample sites A, B, C, E, and F along the Manawatū river. No antimicrobial resistant isolates were isolated from sample site D.

### 6.2 Genetic diversity of ESBL-producing environmental *E. coli*

We used whole genome sequencing to investigate the genetic diversity of 45 of the environmental ESBL-producing *E. coli* (Figure 3, Table S4). Nineteen different sequence types (STs) were identified, with ST131 being the predominant ST (11/45, 24.4%) followed by ST1722 (6/45, 13.3%), ST10 (3/45, 6.7%) and ST7476 (3/45, 6.7%). The predominant ESBL coding gene was *bla*_CTX-M-15_ (28/45, 62.2%, 95% CI: 48.0 – 76.4%) followed by *bla*_CTX-M-27_ (8/45, 17.8%, 95% CI: 6.6 – 29.0%) and *bla*_CTX-M-14_ (7/45, 15.6%, 95% CI: 5.0 – 26.2%). Twenty-one of the isolates had a multidrug resistance genotype. The number of ARGs ranged from 1 to 19, covering ten antibiotic classes: beta-lactams, aminoglycosides, rifamycins, tetracyclines, sulphonamides, trimethoprim, amphenicols, macrolides, phosphonic acids, and lincosamides. However, the resistance genotype was not always concordant with the resistance phenotype. For example, trimethoprim/sulfamethoxazole resistant *E. coli* isolates did not aways have a *dfr* and/or a *sul* gene present.

**Figure 3.**
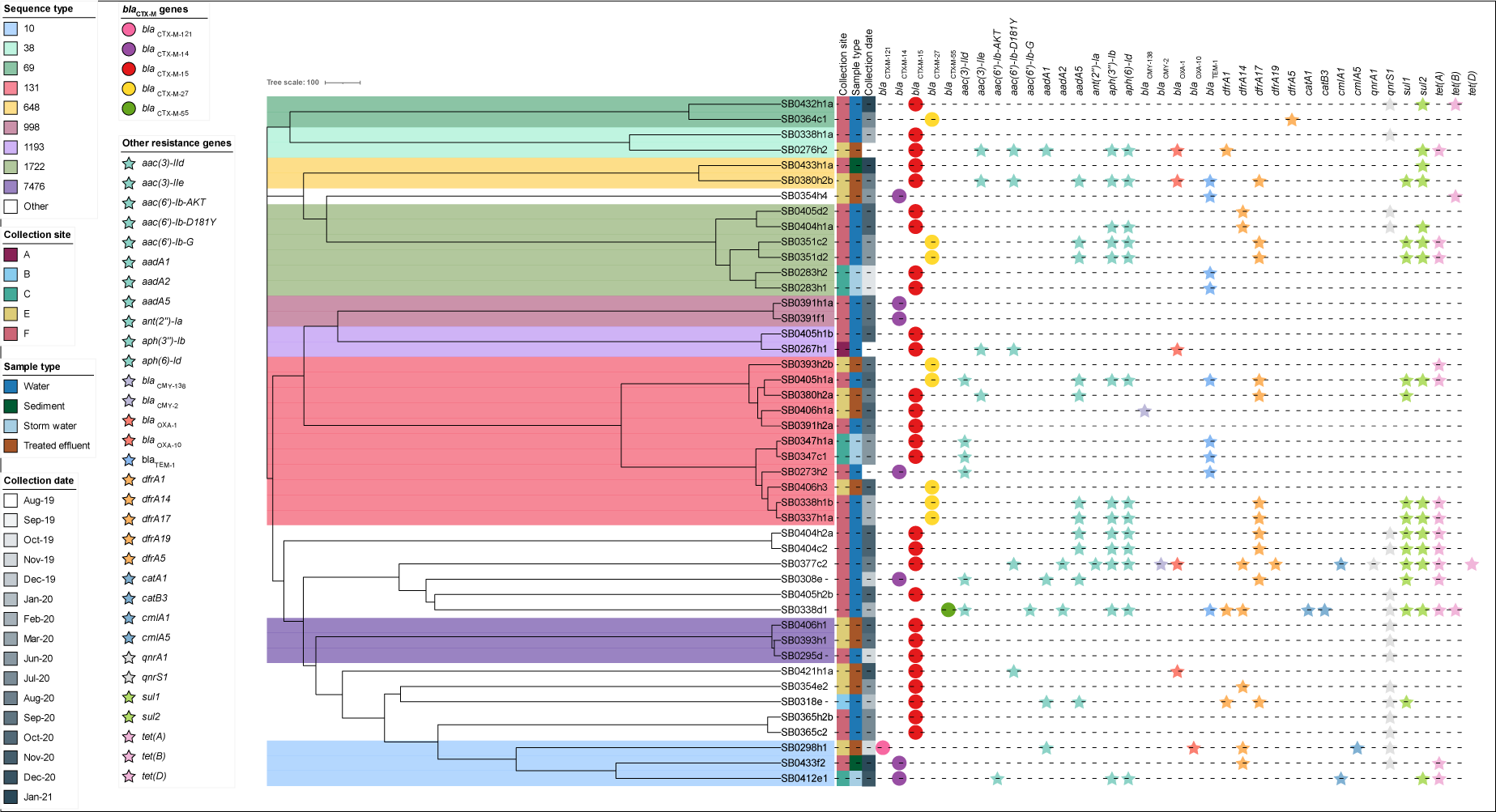
Core-genome MLST neighbour-joining tree, of 45 environmental *E. coli* isolated from river water, sediment, storm water and treated effluent. The tree was generated using 2918 shared alleles identified using chewBBACA.

### 6.3 The genetic relationship of ESBL-producing clinical and environmental *E. coli*

To determine whether there was community spread of ESBL-producing *E. coli* between humans and waterways a comparative genomic analysis was carried out, encompassing 307 human clinical ESBL-producing *E. coli* and the 45 environmental isolates (Table S4). Over the 14 months of sampling there appeared to be no seasonal trend, with ST131 (158/352, 44.9%) being the dominant ST (Figure S2). A cgMLST analysis demonstrated that the environmental isolates were dispersed throughout the phylogeny (Figure 4). There were seven river-only isolate STs (ST156, ST219, ST442, ST542, ST1324, ST1584, and ST2079), two effluent-only STs (ST540 and ST635), and 28 human-only STs. The predominant STs across all the isolates were ST131 (158/352, 44.9%), ST1193 (26/352, 7.4%), ST69 (25/352, 7.1%), ST38 (23/352, 6.5%), ST648 (21/352, 6.0%) and ST998 (15/352, 4.3%). Three STs, ST38, ST131, and ST648, were shared across the three sample types (clinical, effluent and river). Three STs were shared across the clinical and river isolates (ST1193, ST69 and ST998). The predominant ESBL coding genes from the clinical isolates were *bla*_CTX-M-15_ (134/307, 43.6%, 95% CI: 38.1 – 49.1%) and *bla*_CTX-M-27_ (134/307, 43.6%, 95% CI: 38.1 – 49.1%) followed by *bla*_CTX-M-14_ (31/307, 10.1%, 95% CI: 6.7 – 13.5%). The number of ARGs ranged from 1 to 20, covering resistance to ten antimicrobial classes. For the three main STs the predominant ESBL coding gene was *bla*_CTX-M-27_ for ST131, *bla*_CTX-M-15_ for ST69, and *bla*_CTX-M-27_ for ST1193.

**Figure 4.**
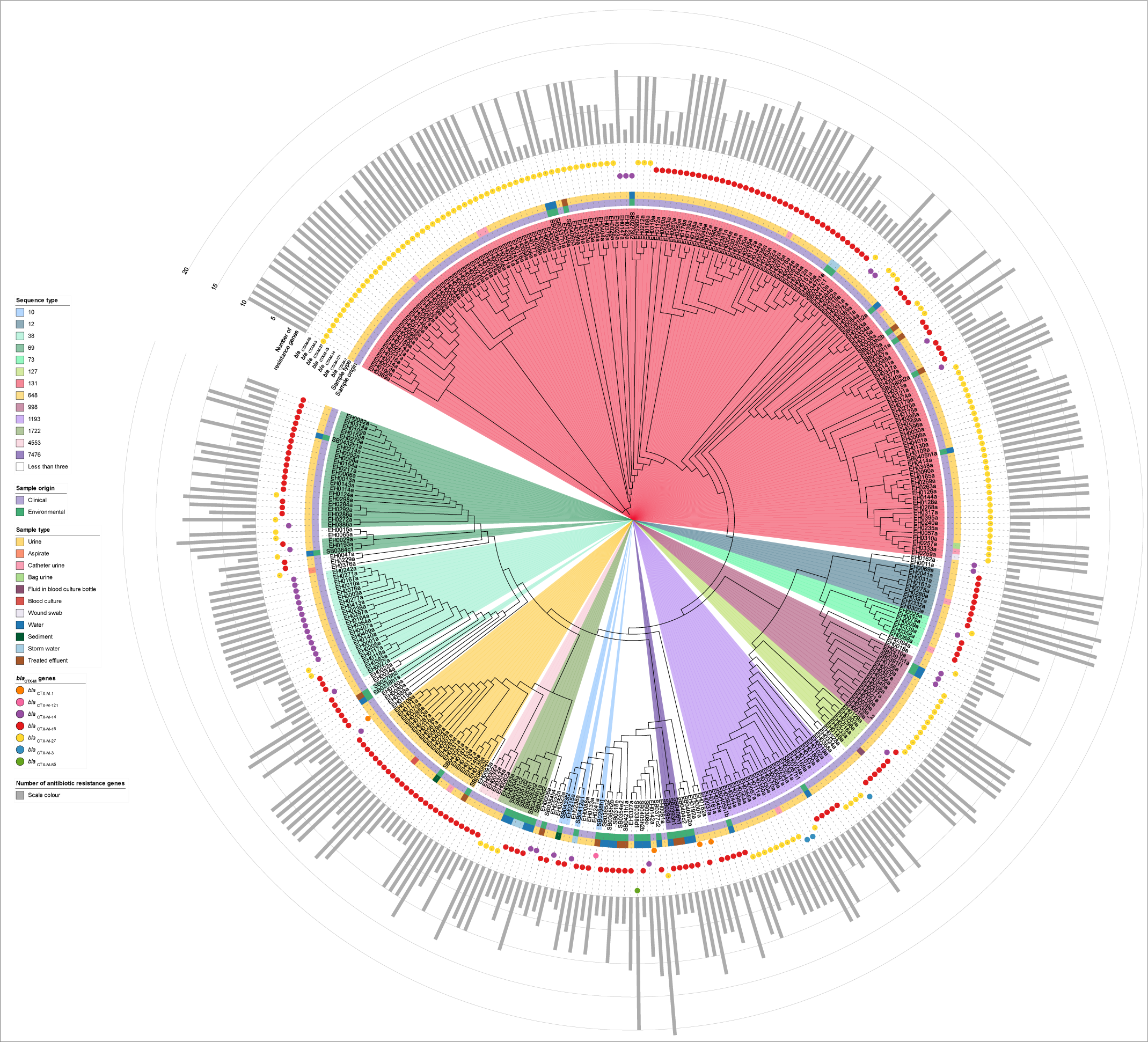
Core-genome MLST neighbour-joining tree of 352 clinical and environmental *E. coli*. The tree was generated using 3,053 shared alleles identified using chewBBACA.

The majority of ST131 isolates fell within clade C (103/158, 65.2%) followed by clade A (51/158, 32.3%), with the 4/158 (2.5%) belonging to clade B being all clinical isolates. The *bla*_CTX-M-27_ was the dominant ESBL coding gene for both the clade A (38/51, 74.5%) and clade C (61/103, 59.2%) isolates. All of the clade C ST131 isolates harboured five point mutations: two in *gyrA*, two in *parC* and one in *parE*; whereas the majority of clade A (44/51, 86.3%) isolates carried two mutations, one each in *gyrA* and *parE*. The ST131 isolates harboured genes typical of ExPEC and UPEC (Table S7), which included the key genes *papA* (1/158, 0.6%), *papC* (77/158, 48.7%), *afaC* (21/158, 13.3%), *kpsM* (146/158, 92.4%), *iutA* (1/158, 0.6%), *chuA* (158/158, 100%), *fyuA* (158/158, 100%), and *sat* (132/158, 83.5%).

We explored the clonal spread of both clinical and environmental ESBL-producing *E. coli* within the three dominant STs (ST131, ST69, and ST1193) by performing a core SNP comparison (Figures 5 and 6, Tables S5, S6 and S7). There was evidence of clonal spread between humans and the environment for ST131 and ST69, but not ST1193. For the ST131 isolates there was less than ten SNPs difference between the environmental ST131 isolates SB0337h1a and SB0338h1b and ten of the clinical isolates, as well as between SB0405h1a and three of the clinical isolates. Two of the clinical strains were received within 14 days of isolating SB0337h1a and SB0338h1b. For the ST69 isolates there was ten SNPs difference between SB0432h1a and three clinical isolates. All four of these environmental strains (SB0337h1a, SB0338h1b, SB0405h1a, SB0432h1a) were isolated from the Manawatū River downstream of the effluent outlet. Although there was no evidence of clonal spread of ST1193 between humans and the environment, the SNP analysis did suggest there was human to human transmission. Isolates EH0056a and EH0097a were isolated from samples collected on the 8^th^ of October 2019 and 19^th^ of November 2019 respectively, with one SNP difference between them, both containing the *bla*_CTX-M-3_ gene, and the same antimicrobial resistant gene profile. There were 4-5 SNPs difference between the isolates EH0087a, EH0111a, and EH0350a. However, EH0111a was received 29 days after EH0087a and EH0350a was received another 11 months later.

**Figure 5.**
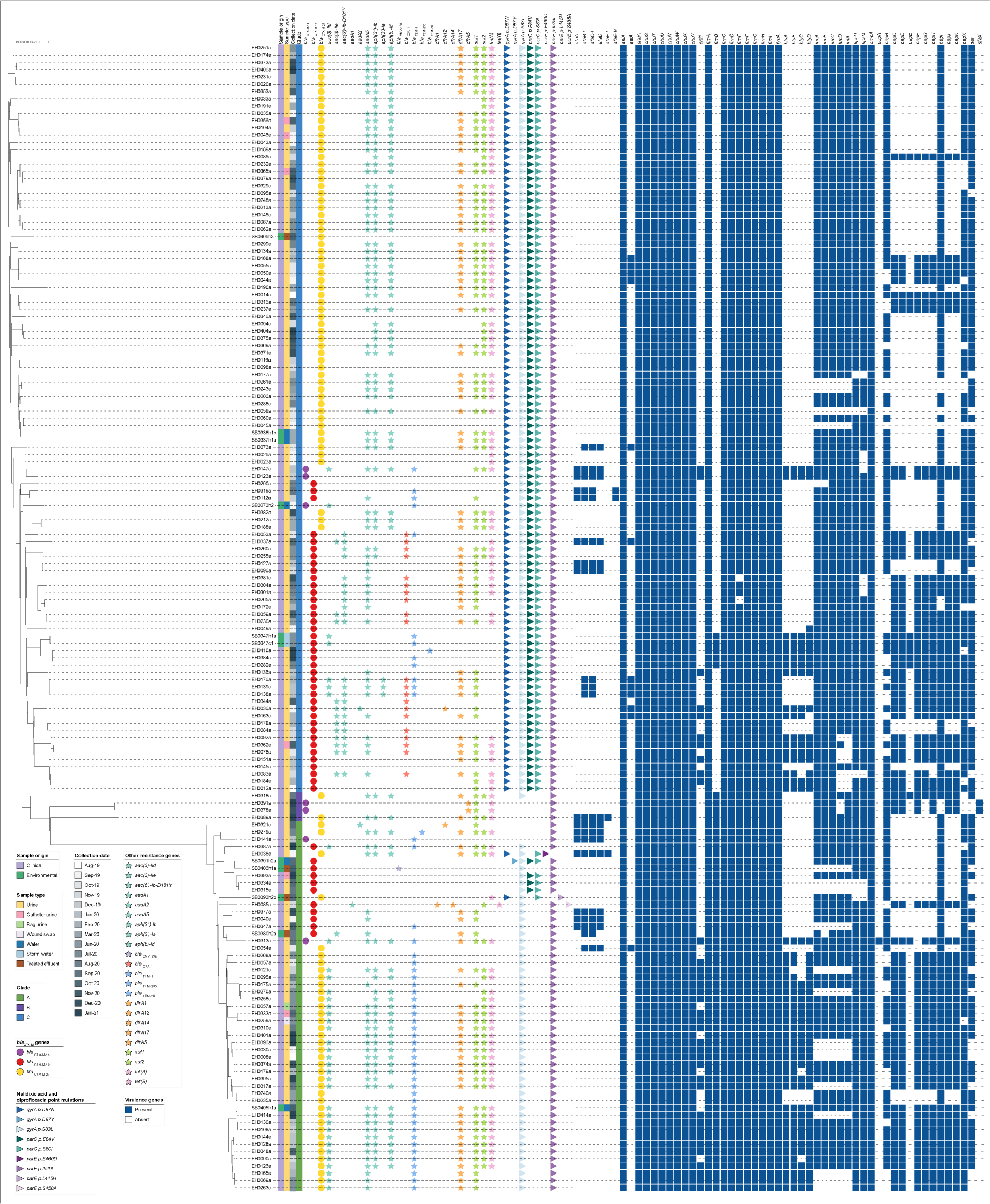
Core SNP phylogeny of clinical and environmental ST131 isolates. Maximum likelihood phylogenetic tree of 159 ST131 *E. coli* produced using 2,370 SNPs with EH0395a used as the reference genome. The tree was constructed with FastTree using a maximum-likelihood GTR model and visualised in iTOL.

**Figure 6.**
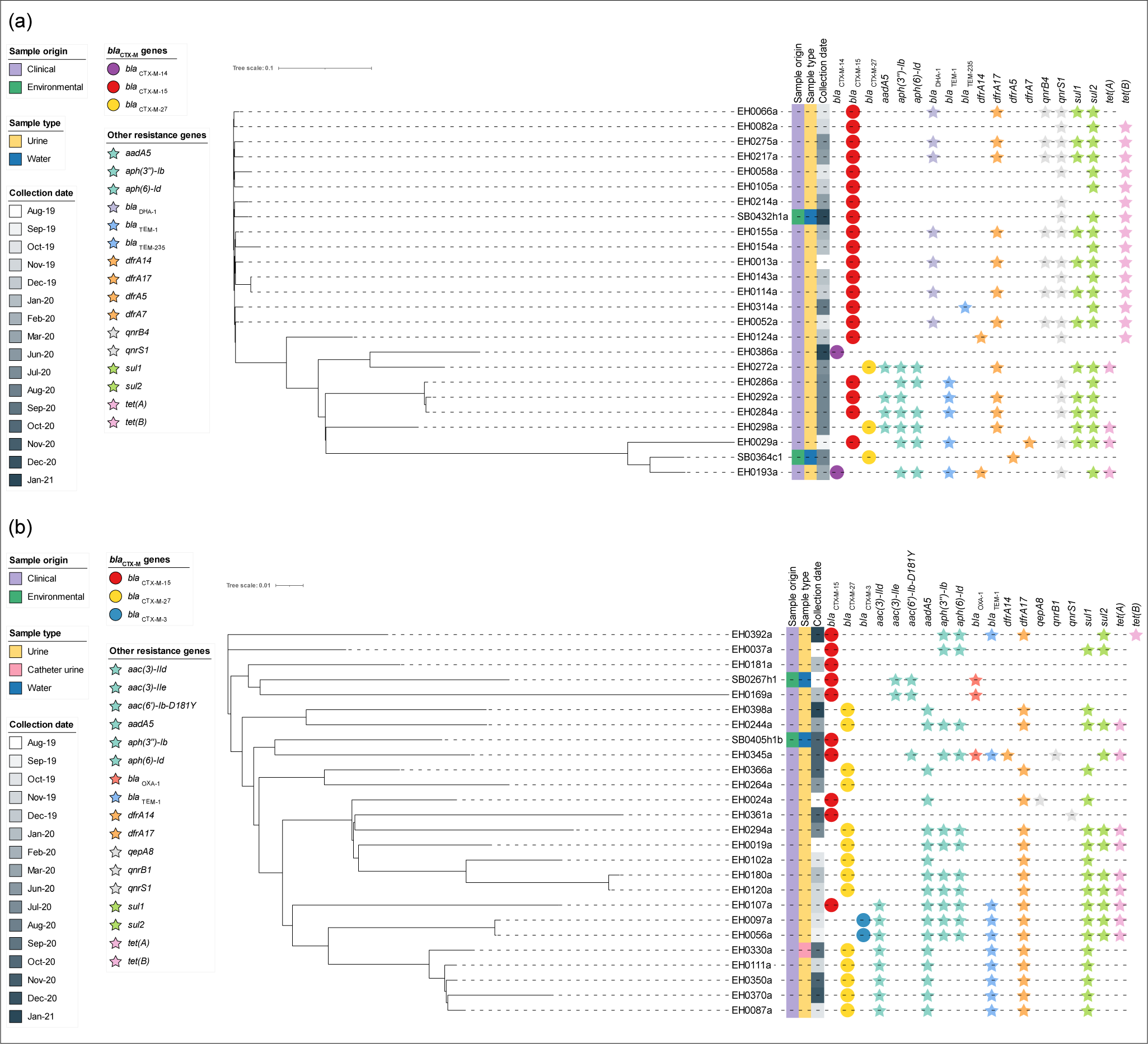
Core SNP phylogeny of ST69 and ST1193 clinical and environmental *E. coli* isolates. a) Maximum likelihood phylogenetic tree of 25 ST69 *E. coli* produced using 630 SNPs with EH0143a used as the reference genome. b) Maximum likelihood phylogenetic tree of 26 ST1193 isolates produced using 726 SNPs with EH0294a used as the reference genome. The tree was constructed with FastTree using a maximum-likelihood GTR model and visualised in iTOL.

### 6.4 Plasmid analysis

The plasmidome genes across all 352 genomes were compared (Figure 7. There was some clustering by ST, plasmid Inc type and *bla*_CTX-M_ variant. There were also clusters with multiple STs, but the same Inc type and *bla*_CTX-M_ variant. To determine the probable origin of the *bla*_CTX-M_ genes, a plasmidome analysis was carried out, in which 144 contigs with *bla*_CTX-M_ genes were identified as originating from plasmids, of which 11 carried a *bla*_CTX-M_ gene and plasmid replicon on the same contig. Three different Inc types (IncFII, IncI and IncB/O/K/Z) were associated with these 11 contigs carrying *bla*_CTX-M_ genes.

**Figure 7.**
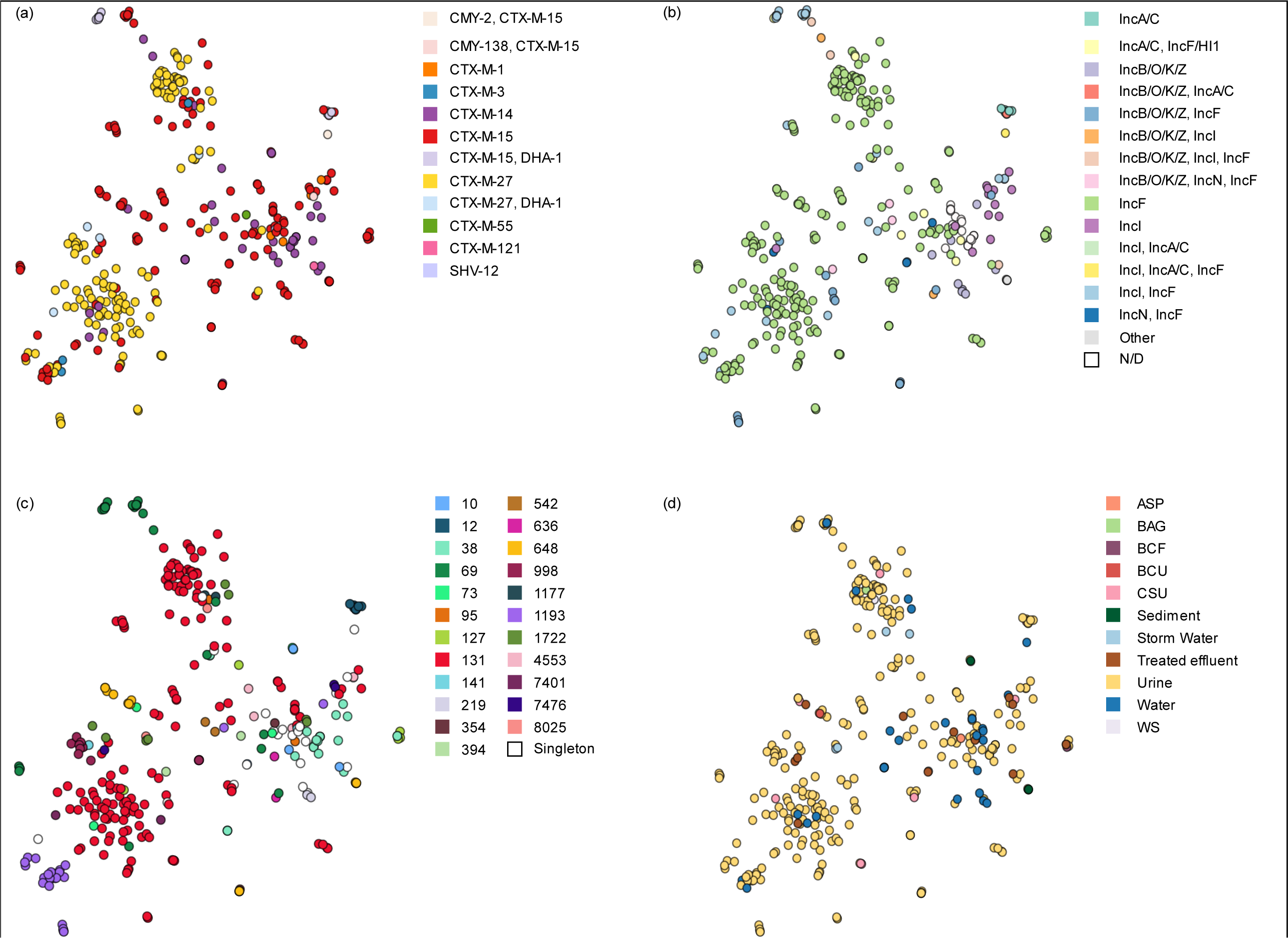
Plasmidome network of the plasmid contigs from 352 clinical and environmental *E. coli*. Each circle denotes the concatenated plasmid contigs from one isolate and is coloured by a) sequence type, b) Inc type, c) ESBL and AmpC variant and d) source of isolate. ASP: aspirate, BAG: bag urine, BCF: aspirate fluid (generally blood) in blood culture bottle, BCU: blood culture, CSU: catheter urine, WS: wound swab.

## 8. Discussion

Our study found that ESBL-producing *E. coli* were frequently isolated from an Aotearoa New Zealand waterway that passes through an urban environment. These bacteria were isolated from at least one sampling site during 12 out of the 13 time-points. Three of the four sampling sites were associated with an urban land-use, while one site was associated with agricultural land-use. The site with the highest prevalence of ESBL-producing *E. coli* was located downstream of a treated effluent outlet (site F). This is not surprising as it has previously been shown that wastewater treatment is not completely effective at removing antimicrobial resistant bacteria [59–62], and studies have found a higher prevalence of ESBL-producing *E. coli* and other types of resistant *E. coli* immediately downstream of effluent outlets [32, 63–65]. A higher prevalence of antimicrobial resistant bacteria and their genes has also been detected during periods associated with higher rainfall [66–68]. One reason for this increase could be sewage overflow, which has been shown to be the dominant contributor to the bacterial community in a river downstream of a wastewater treatment plant after high rainfall events [69]. In our study ESBL-producing *E. coli* were isolated throughout the year during periods of low and high rainfall. These bacteria were also isolated from sites upstream of the wastewater treatment plant. In Aotearoa New Zealand, it has previously been established that there are antimicrobial resistant *E. coli* present in freshwater environments [30, 35, 70, 71]. Studies suggest that that the recreational use of contaminated waters may also be an exposure route for ESBL-producing *E. coli* associated infections in humans [21, 25]. Although, the carriage of ESBL-producing *E. coli* has not been found to be higher in recreational water users, previous studies have identified recreational water use as a risk factor for ESBL-producing *E. coli* associated infections [26, 28].

The most common ESBL coding gene variant among *E. coli* isolated from waterways differs between studies. Our study found that *bla*_CTX-M-15_ was the dominant gene variant for the treated effluent, and water, which concurs with studies carried out in Europe [32, 33, 72]. In contrast, a study carried out in Brazil found *bla*_CTX-M-2_ was the dominant gene variant for ESBL-producing *E. coli* isolated from a waterway downstream of a wastewater treatment plant, whereas *bla*_CTX-M-8_ was the main ESBL coding gene associated with *E. coli* isolated from the wastewater plant [73]. Other *bla*_CTX-M_ variants commonly harboured by ESBL-producing *E. coli* from freshwater include *bla*_CTX-M-1_, *bla*_CTX-M-14,_ *bla*_CTX-M-27_ [74–78].

The most frequently detected ESBL gene types amongst the clinical isolates were *bla*_CTX-M-27_ and *bla*_CTX-M-15,_ with both having a proportion of 134/307 (43.6%). These findings are similar to a survey recently undertaken in Aotearoa New Zealand, which found 68/158 (43.0%) human clinical ESBL-producing *E. coli* isolates carried the *bla*_CTX-M-15_ and 64/158 (40.5%) carried the *bla*_CTX-M-27_ gene [6]. A regional survey also found a similar proportion of ESBL-producing *E. coli* carrying *bla*_CTX-M-27_ compared with *bla*_CTX-M-15_, 18/65 (27.7%) and 14/65 (21.5%) respectively [13].

In concordance with other studies [72], other ARGs conferring resistance to a range of other antibiotic classes including aminoglycosides, trimethoprim, sulphonamides, tetracyclines, fluoroquinolones, phenicols, and phosphonic antibiotics were present across both the clinical and environmental isolates. Similar plasmid types were also shared across both the clinical and environmental isolates, with IncF being the most common plasmid type. IncF plasmids frequently harbour *bla*_CTX-M_ genes [79]. The close clustering of isolates by their plasmidome across some STs in our study suggests horizontal gene transfer may have occurred between strains. Long-read sequencing of plasmids would be needed to confirm which plasmid Inc types harboured the ESBL coding genes.

Both treated effluent and water samples from the river contained a diverse range of *E. coli* sequence types, in agreement with previous studies [32, 33, 62]. ST131, which is the main lineage associated with UTIs and blood infections, was the dominant type for the effluent, river samples, and clinical isolates. In Aotearoa NZ, ST131 remains the most prominent ST found in clinical specimens as reported by previous surveys [6, 13]. In our study, the majority of ST131 clinical and environmental isolates fell within clade C. In contrast, a recent wastewater surveillance study in Canada found that clade A was the dominant ST131 clade with only 8.6% of strains belonging to clade C [80]. In our study, all the clade C isolates had a fluroquinolone resistance genotype. Elevated levels of fluoroquinolone resistance are generally associated with mutations in both *gyrA* and *parC* [2, 81], and are commonly associated with the C2 clade but is reported to be rare in clade A strains, as was also found in our study [82]. However, a recent study found 72.7% of ST131 clade A strains isolated from wastewater in Canada were resistant to ciprofloxacin [80].

The other dominant ST from the water samples was ST1722. This ST has previously been isolated from humans, sewage, livestock, birds, companion animals and waterways (https://enterobase.warwick.ac.uk/, accessed 13 February 2024). Interestingly, in our study ST1722 was only isolated from water. ST38 and ST648 were isolated from humans, sewage, and river samples. ST38 and ST648 are often associated with blood infections and UTIs in humans and have frequently been isolated from waterways and sewage [32, 33, 62, 83–85]. A previous study conducted in Sweden also isolated ST38 *E. coli* from water samples downstream of a treated effluent outlet. Other studies that have examined freshwater for the presence of ESBL-producing *E. coli* have found that ST949 and ST10 dominate [33, 72]. ST949 was not isolated from the river and ST10 was isolated from the sediment once during our study.

In this study we found that ESBL-producing *E. coli* found in an Aotearoa NZ waterway were genetically similar to those isolated from human clinical infections, where the difference in the number of SNPs was less than or equal to 10 SNPs between several human and river isolates. However, our epidemiological data only supported the recent transmission between humans to the local river within one set of isolates, where the dates of isolation of both the human and the river isolates were within 14 days. The close clustering of multiple human clinical isolates also suggests that there was human-to-human transfer or contact with the same source of ESBL-producing *E. coli* in the community. A limitation to our study is that we only sampled from one geographical area and within this area the treated effluent and river collection occurred once a month, whereas our clinical isolates were collected weekly. This may have reduced our ability to detect more ESBL-producing *E. coli* from the treated effluent and river that were genetically similar to the clinical isolates.

Previous studies have also indicated that there is the spread of ESBL-producing *E. coli* from humans to freshwater (or *vice versa*) [30, 32, 72]. Fagerström, Mölling [32] compared ESBL-producing *E. coli* UTI isolates to those sourced from fresh water using a cgMLST approach and found that some isolates had less than ten allele differences, suggesting the sharing of strains between humans and the environment. A study conducted in Germany found that ST949 *E. coli* isolates collected from swimming and bathing sites were closely related to human clinical ST949 isolates from Aotearoa New Zealand and Sweden, although the difference in the number of SNPs or allele changes between isolates was not stated [72]. The most likely source of human-associated ESBL producing *E. coli* is through the disposal of treated effluent into our waterways [86].

In conclusion, this study found that ESBL-producing *E. coli* are present in water, sediment, stormwater, and treated effluent samples collected along the Manawatū River. It was shown that while treated effluent is a source of antimicrobial resistant *E. coli*, these resistant bacteria were also present in the Manawatū River upstream of the treated effluent outflow. There was some evidence for the sharing of genetically related ESBL-producing *E. coli* between clinical and environmental sources. The study was limited by the number of isolates collected and therefore sequenced. More frequent sampling would provide a clearer picture of the genetic relatedness between environmental and human ESBL-producing *E. coli* isolates. Our findings emphasise the importance of including an environmental component to antimicrobial resistance surveillance.

## 9. Author statements

### 9.1 Author contributions

H.G: Methodology, formal analysis, investigation, funding acquisition, writing – original draft. P.J.B: Genomic analysis, investigation, supervision, funding acquisition, writing – review and editing A. M: Study conceptualisation, methodology, investigation, supervision, funding acquisition, writing – review and editing. L.R: Methodology, resources, investigation, writing – review and editing. A.F: Data curation, investigation, writing – review and editing. R.A: Methodology, investigation, writing – review and editing. S.B: Study conceptualisation, methodology, investigation, funding acquisition, supervision, formal analysis, writing – original draft.

### 9.4 Ethical approval

This project has been evaluated by peer review and judged to be low risk (ethics notification number: 4000021252). Consequently, it has not been reviewed by one of Massey University’s Human Ethics Committees. This study used bacterial strains, which were isolated from human samples. No biological material of human origin was obtained for this study and the origin of samples was anonymised.

## 9.5 Acknowledgements

We thank Medlab Central for supplying the clinical Enterobacterales isolates and Massey Genome Services for their assistance with the whole genome sequencing. We wish to acknowledge the use of New Zealand eScience Infrastructure (NeSI) high-performance computing facilities, as part of this research.

