## Supplementary figures and images for "Genomic epidemiology of ESBL-producing *Escherichia coli* from humans and an Aotearoa New Zealand river"

### Supplemental Figure 1

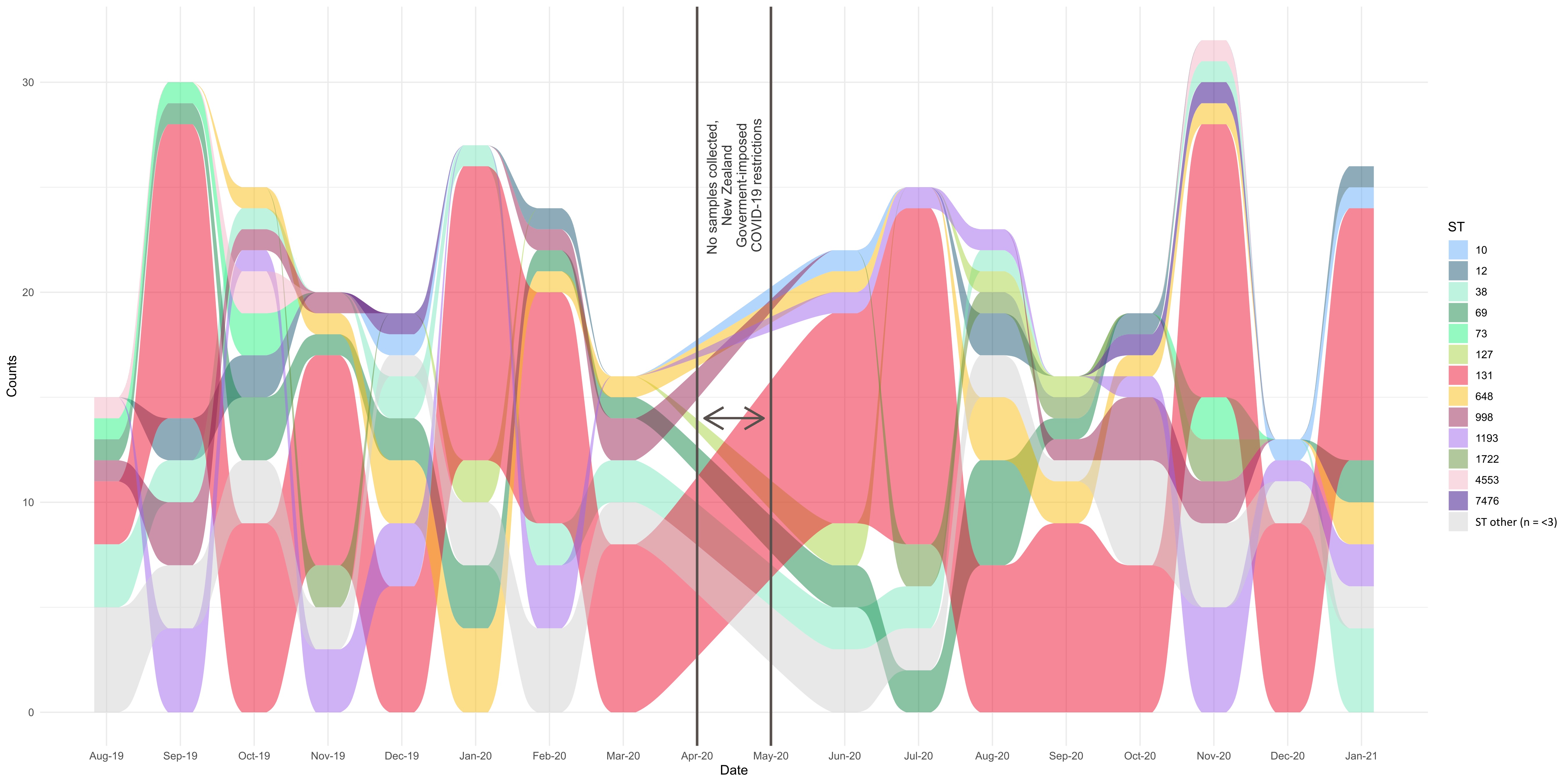
